# Deep learning-based proteomics enables accurate classification of bulk and single-cell samples

**DOI:** 10.1101/2024.02.03.578734

**Authors:** Karl K. Krull, Arlene Kühn, Julia Höhn, Titus J. Brinker, Jeroen Krijgsveld

## Abstract

Proteins are the main drivers of cell function and disease, making their analysis a powerful technique to characterize determinants of cell identity and to identify biomarkers. Current proteomic technology has the breadth to profile thousands of proteins and even the sensitivity to access single cells, however limitations in throughput restrict its application, e.g. not allowing classification of samples according to biological or clinical status in large sample cohorts. Therefore, we developed a deep learning-based approach for the analysis of mass spectrometric (MS) data, assigning proteomic profiles to sample identity. Specifically, we designed an architecture referred to as Proformer, and show that it is superior to convolutional neural network-driven architectures, is explainable, and demonstrates robustness towards batch-effects. Based on its tabular approach, we highlight the integration of all four dimensions of proteomic measurements (retention time, mass-to-charge, intensity and ion mobility), and demonstrate enhanced sample discrimination involving a treatment with IFN-γ, despite its subtle effect on the cell’s proteome. In addition, the Proformer is not restricted to proteomic depth, and can classify cells by cell type and their differentiation status even using single-cell proteomic data. Collectively, this work presents a novel deep learning-based model for rapid classification of proteomic data, with important future implications to enhance patient stratification, early detection and single-cell analysis.

## Introduction

Proteins control nearly all biochemical processes in cells, and their expression is dynamically regulated to meet cellular demands during development or in response to external cues. The collective set of proteins that make up the proteome can therefore be regarded as a close representation of the cell′s functional phenotype, and its investigation gives detailed insight into the role of proteins in basic cellular processes and in disease (1). Most proteomic studies use peptides obtained by proteolytic digestion as a representation of the cellular protein composition as they are more amenable for detailed characterization by proteomic techniques (2, 3). This usually entails peptide separation according to their retention time in liquid chromatographic (LC) systems and in mass spectrometers (MS) that determine their mass-to-charge (m/z) ratio, while integration of m/z intensities over time is used for peptide quantification. In a second step, peptides are fragmented to generate information that is used to infer protein identity, and this strategy has been used in a vast number of studies to understand proteome adaptation in multiple biological and clinical settings. However, instead of going through the process of protein identification, peptide features in terms of retention time, m/z and intensity can be used to define a sample’s peptide profile that is characteristic of its proteomic make-up, and there are increasing efforts to utilize this information in deep-learning (DL) methods that classify samples based on their LC-MS fingerprint (4). Such an approach can significantly increase analysis speed and data interpretation, which in turn will enhance sample throughput, and would be of particular relevance in a clinical context for classification of patients or disease states requiring fast turn-around times. As the most prominent example, convolutional neural networks (CNNs) have been applied to classify proteomic samples by converting their peptide profile into a heatmap-like image, distinguishing conditions based on their spatial peak patterns (4-8). To this end, CNNs rely on binning data of proximate signals to achieve a uniform grid input, and to employ pre-trained networks (4), which aligns individual samples that suffer from variabilities in the measurement (6). Nevertheless, this requires complex pre-processing, and it is unclear to what extent binning affects the data and may cause loss of information. Moreover, CNNs are computationally heavy, prone to overfitting and limited to a fixed number of feature dimensions due to its image input, making it inaccessible to recent advances in MS instrumentation that implement ion mobility as a fourth separation layer (9). As an innovative alternative, transformer models offer simpler architectures based on a self-attention mechanism that learn complex relationships at low computational costs and higher resistance to overfitting (10), while they have already been implemented in various fields, e.g. for natural language processing (11), image tasks (12) and tabular data (13).

Here, we introduce the Proformer (Proteome Transformer), a transformer-based DL model, built to classify proteomic samples accurately and with high robustness using their tabular peptide profile. The flexible architecture of the Proformer allows the implementation of any desired number of feature dimensions derived from MS data, including ion mobility, and preserves peptide information without prior binning. Using this approach, we achieve higher accuracies than CNNs for a large, in-house generated data set involving a treatment with IFN-γ, despite its mild effect on the cell’s proteome. We show that transformer-based architectures are less vulnerable to batch effects and specifically investigate the effect of quantitative differences in the experimental conditions on the achievable classification accuracy. To this end, we use several recently published data sets and demonstrate that DL models have the capability to discriminate samples even at the single-cell level, qualifying it for broader application in the future.

## Results

Current widespread proteomic measurement lacks throughput while generating high-dimensional data with thousands of identified peptides in a limited number of samples, restricting its accuracy and making it susceptible to overfitting in deep neural networks. To overcome this, we developed the Proformer model, a proteome classifier that is based on a transformer approach using a self-attention mechanism. Since its recent successes in other application fields, we were interested in evaluating this type of DL architecture in the challenging adaption to proteomic samples, and compared it to formerly used image-based CNN approaches. To this end, we critically assessed their performances in the classification of HeLa cell samples that were treated with Interferon gamma (IFN-γ) (**Fig. 1A**), an extensively described cytokine that specifically alters the expression of its target proteins, but only results in mild effects in the proteome of non-immunogenic cells (14, 15).

**Figure 1:**
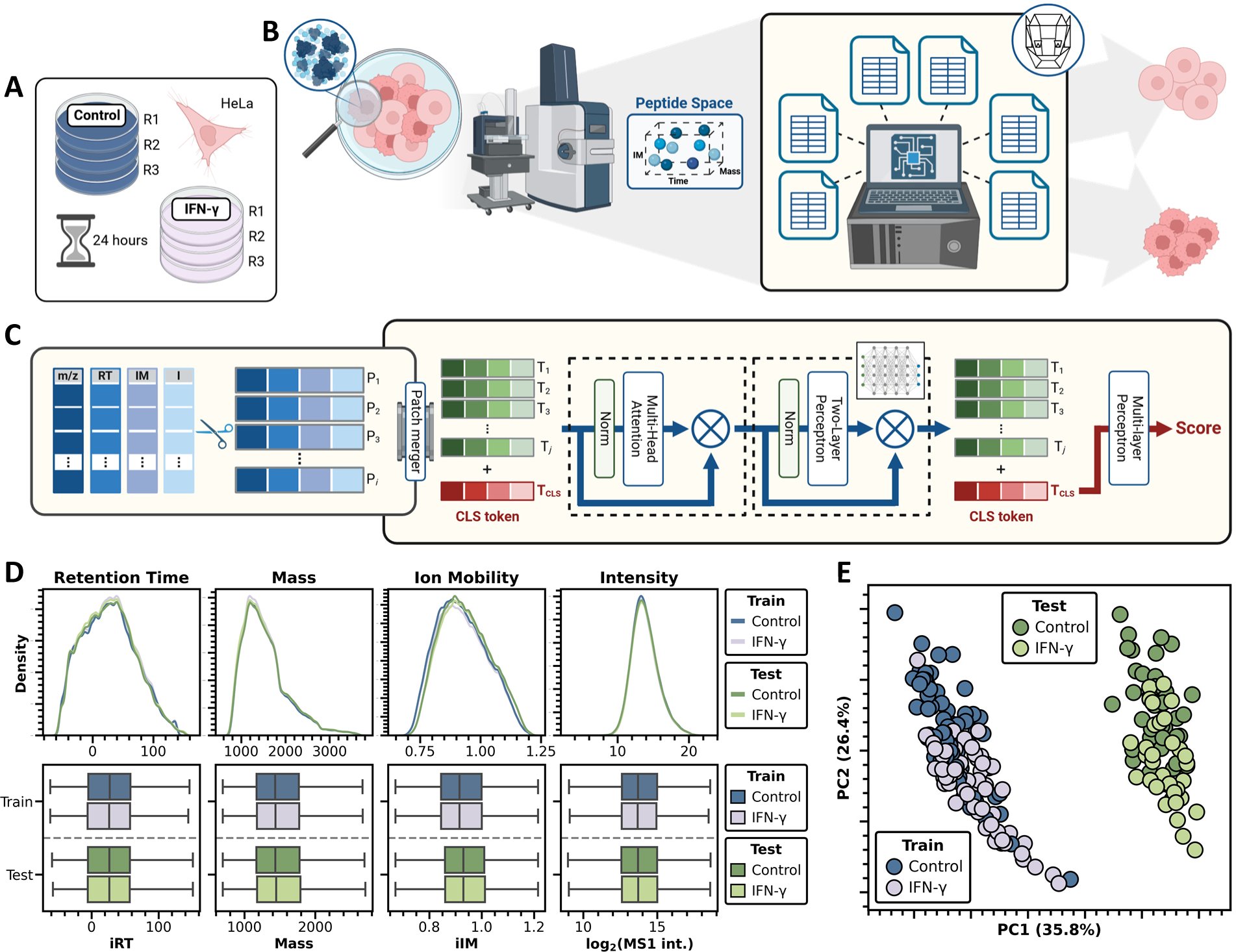
DL-assisted characterization of proteomic samples using Proformer. (A) Data set generation for training and testing of the DL model. Control (dark blue) and IFN-γ-treated (lavender) HeLa cells (three individual cultures R1 – R3, respectively) were aimed to be distinguished after 24 hours of incubation. (B) Conceptual design of DL-assisted classification pipeline: proteomic profiles of samples are defined by their peptide space acquired in a tabular format that is used by the Proformer model to discriminate profiles of two groups. (C) Structure of transformer architecture. Tensors of dimensional peptide data are extracted and reduced to j tokens using a patch merger. For the classification task, a generated CLS token is added. The input is passed to N Encoder blocks and then to the classifier, which is a Single-layer Perceptron. Each Encoder block consists of a Multi-head Attention, followed by a Two-layer Perceptron and normalization layer in between. The CLS token is passed to the classifier, which determines the score based on the input. (D) Distribution of the recorded training (dark blue and lavender) and test data set (greens) along the four acquired dimensions: indexed retention time, mass, indexed ion mobility and MS1 intensity. Top panels: kernel analysis. Bottom panels: boxplot analysis. (E) Peptide-level principal component analysis (PCA) of the training data set (blue: control; lavender: IFN-γ-treated; samples n = 142) and test data set (green: control; light green: IFN-γ-treated; samples n = 71). Explained variance ratios are indicated at both axis.

We generated the training dataset by injecting 100-ng peptide samples of control and IFN-γ-treated HeLa cells (142 total) on a timsTOF Pro instrument utilizing a conventional diaPASEF methodology (**Fig. S1A**) to acquire MS1 and MS2 peptide information. Since we were specifically interested in assessing the batch effect of temporally distant experiments, injections of the test dataset (71 total) were performed six months apart under fixed measurement conditions. Among approx. 145,000 identified peptides and 9,500 proteins per injection in the training set (135,000 peptides and 9,100 proteins per injection for test data) (**Fig. S1B** and **C**), we observed excellent data completeness (**Fig. S1E** and **F**), demonstrating the high quality of the obtained datasets. We extracted the indexed retention time (iR_t_) and ion mobility (iIM), mass-over-charge (m/z) and intensity features of the identified peptides in a DL-based classification task, aiming to discriminate treated from control samples based on their MS1 profile (**Fig. 1B**). Utilizing a transformer approach, we successfully integrated all four feature dimensions, benefitting from its versatile input variables, whereas the conventional CNN pipeline is restricted to only three dimensions due to its image conversion (**Fig. S2**). Conceptually, the obtained peptide matrix is passed to a patch merger that reduces dimensionality to a number of tokens, while it preserves the number of introduced features (**Fig. 1C**). The transformer proceeds with an architecture of transformer layers and a classifier, in which each transformer block consists of a multi-head attention followed by a multi-layer perceptron module, and an adjacent drop-out and normalization layer, respectively. In this way, the classification task only demands an encoder block as transformer layer, while the concatenated CLS token eventually gives rise to a label prediction for a given input (**Fig. 1C**).

To assure that the desired classification will be solely based on quantitative differences specifically in peptides that are responsive to the treatment, we verified the technical equality of the obtained datasets. In this regard, extracted features from training and test set showed almost identical distributions, highlighting the consistency of the measurement, while even the control and IFN-γ treatment conditions displayed minor differences (**Fig. 1D**). On the other hand, quantitative differences were indeed evident in the data sets, leading to the formation of two cohesive cluster in peptide principal component analysis (PCA), which are divided in PC1 according to their time of acquisition as intended (**Fig. 1E**). Importantly, treated and non-treated samples separated along PC2, although not leading to distinct clusters, likely as a result of modest IFN-γ-induced effects in the proteome. Consequently, we designed a particular difficult task, in which the DL algorithm needs to transfer relations of treatment effects from one batch to another, generalizing the classification problem.

Having ensured the integrity of our data, we applied the training set to tune different classification models using the transformer architecture (**Fig. 1C**) or the CNN image pipeline (**Fig. S2**) in a five-fold cross validation approach, and determined their performance by the test dataset (**Supplementary data 1**). Remarkably, our 3D-Proformer model (based on retention time, m/z and intensity) showed accurate classification of experimental conditions, resulting in correct grouping of 83.3% of the samples, compared to up to only 79.9% in the CNN approaches (**Fig. 2A**).

**Figure 2:**
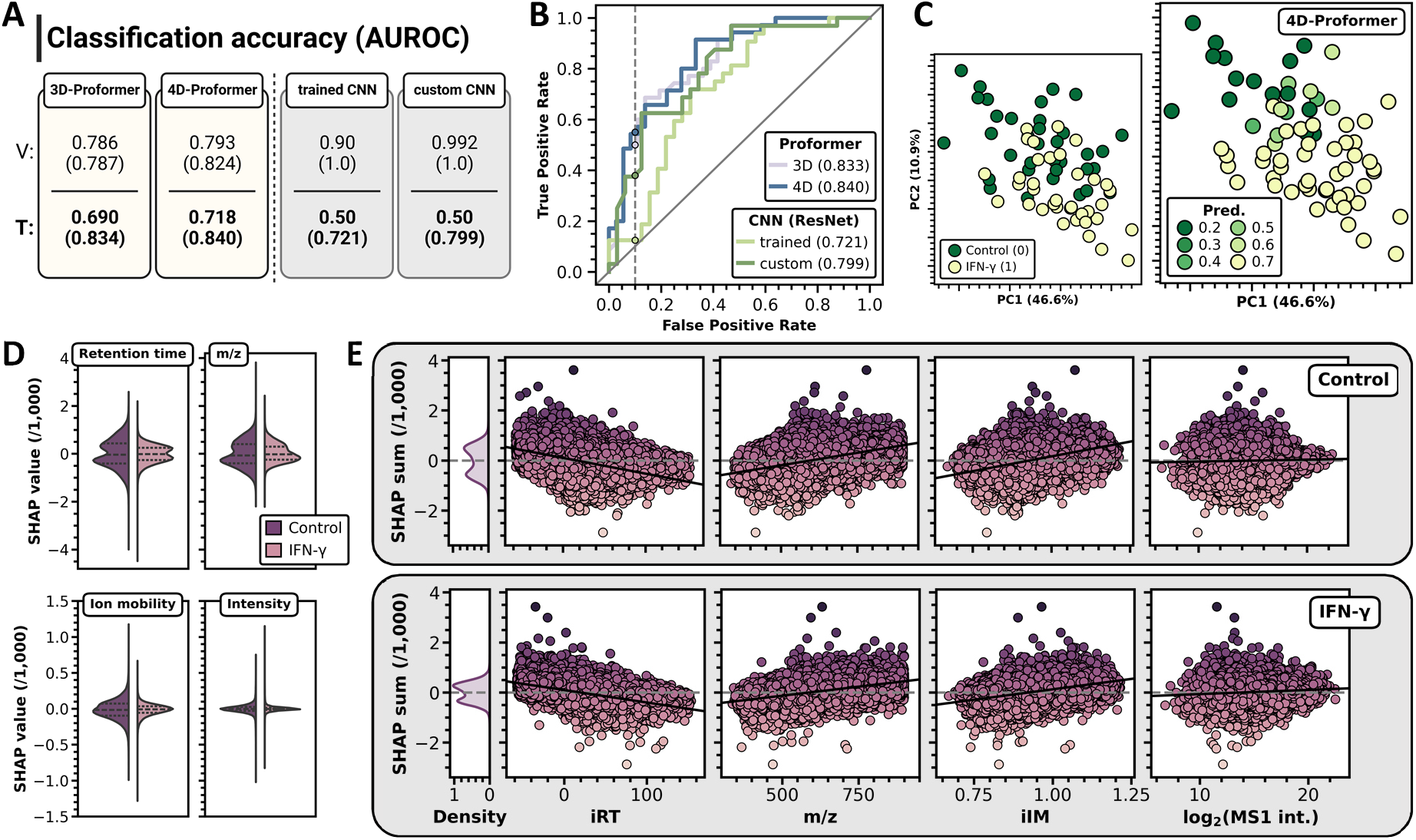
Proformer enables accurate classification of time-lapsed proteomic samples. (A) Accuracy (label cutoff: 0.5) and AUROC values (in parentheses) for the classification of validation and test set using the Proformer (yellow: based on three and four feature dimensions) or the CNN image approach (grey: pre-trained and newly trained ResNet_v2_50). (B) Receiver operating characteristics (ROC) over Proformer (lavender and blue) and CNN image (light and dark green) models. Specified points indicate the observed sensitivity (true positive rate) at a specificity (1 – false positive rate) cutoff at 90% (dashed line). Diagonal line represents a random classifier. Areas under ROC (AUROC) are indicated in parentheses. (C) Peptide-level PCA of the test set showing quantitative differences of control (dark green, training label 0) and IFN-γ-treated samples (yellow, training label 1) (left panel), and predicted labels of the 4D-Proformer model (right panel). Explained variance ratios are indicated at the axes. (D) SHAP (Shapley additive explanation) value distributions of respective feature dimensions for control and IFN-γ-treated samples (divided violins) in the 4D-Proformer model. Dashed lines in the violins represent median, first and third quartile of the dataset. (E) Distributions of SHAP value sums of control and IFN-γ samples showing learned dependencies on the respective feature dimensions (black line: linear regression by Pearson correlation) and beneficial/detrimental combinatorics of features for peptide data points.

Notably, the inclusion of IM as a fourth feature dimension further improve accuracy to 84.0% (4D-Proformer). This was also reflected by the receiver operating characteristic (ROC), where the Proformer approaches showed superior sensitivities, e.g. at a specificity cut-off of 90% (**Fig. 2B**). As an interesting notion, performance of the CNNs was strongly dependent on their initial weighting before training, with the custom-trained network achieving significantly higher AUROC (0.799) than the pre-trained network (0.721) in the test set (**Fig. 2A** and **B**). Moreover, we observed a notable discrepancy in the CNN results, which showed a drop in performance of around 20 – 28% from the validation to the test set, while the Proformer model reached equivalent accuracies in both data (**Fig. 2A** and **B**). Since validation is part of the training procedure, these results suggest that the tabular approach in the Proformer model is much better at dealing with the batch differences between validation and test sets. This might be due to the signal binning in the CNNs that groups peptide peaks with similar retention time and m/z values, and might therefore be more prone to small shifts that lead to altered bins and images (**Fig. S2**).

To improve our understanding of the 4D-Proformer, we analyzed the contribution of the respective properties of the input data on the output of the model. Specifically, we located samples based on their predicted label (0: control, 1: IFN-γ-treated) in a PCA analysis and found misclassified samples exclusively in the overlapping region of the two experimental conditions, while distant samples showed a correct assignment (**Fig. 2C**). This observation shows that the model’s decisions are indeed determined by the peptide intensity profile of control and IFN-γ-treated samples, and emphasizes the role of quantitative differences for the difficulty of the classification task. Next, we analyzed the test data using the SHAP (SHapley Additive exPlanation) value system (16), a method for interpreting the output of machine learning models, in which the feature dimensions are assigned an importance metric to assess their contribution to the prediction of the model (positive values: positive impact; negative values: negative impact). Contrary to our expectation, retention time and m/z dimensions constituted the dominant factors considered by the Proformer, while intensity values showed significantly less impact (**Fig. 2D**). This observation might be because retention time and m/z can be determined with higher precision, which the model therefore may consider more reliable. In this sense, a higher sample number might augment the contribution of peptide intensity. Nevertheless, the analysis also showed a notable contribution of the ion mobility feature, justifying its implementation in the Proformer approach by possibly explaining its positive effect on the classification accuracy. Further analysis of feature-specific SHAP values revealed a positive impact of peptides with low retention time but high m/z and negative impact of those with high retention time but low m/z (**Fig. S3A**). While these aspects compensated each other for most peptides, global dependencies were preserved even when SHAP values per peptide were summed (indicated by linear regressions) (**Fig. 2E**), as peptides with the most pronounced impact still followed these trends. Although we have not investigated this phenomenon in detail, we observed that quantitative variation of individual peptides showed a similar but less distinct dependence on their feature dimensions (**Fig. S3B**). However, it cannot be excluded that there is also an explanation by the underlying cellular effect that drives the model to focus specifically on this relationship.

Collectively, these data demonstrate that the Proformer model is a superior alternative to CNN image pipelines, enabling accurate classification of proteomic samples based on their peptide profile, which are resistant to secondary influences such as batch effects, and allow the implementation of additional feature dimensions during peptide acquisition. Furthermore, we emphasized the importance of methods that facilitate the explanation of DL models and presented SHAP values as a valuable resource to elaborate the rationale of the Proformer approach.

### DL-assisted classification of single cells enabled by the Proformer

Having constructed and successfully tested our Proformer model for LC-MS data of conventional bulk proteome samples, we aimed to extend its applicability to the classification of single cells. The emerging field of single-cell proteomics (SCP) usually involves the measurement of several dozens of samples to investigate the statistical variation of the underlying biological process at stake, making it ideal to provide DL-based models with sufficient training material. With this in mind, we aimed to evaluate the performance of our transformer pipeline and used three recently published single-cell data sets (**Table 1**) to investigate specifically the role of quantitative similarities between the experimental conditions. In this sense, the selected studies varied mainly in the extent of their experimental differences, ranging from comparing distinct cells types (low overlap in tSNE) to subtle biological treatment (high overlap in tSNE, see **Fig. 3**).

**Table 1:** Overview of label-free single-cell studies used for Proformer-based classification.

| No. | Ref. | Resource | Features | Samples:<br>train / test | Peptides<br>per sample | Experiment | tSNE<br>overlap | AUROC<br>test set |
| --- | --- | --- | --- | --- | --- | --- | --- | --- |
| 1 | (17) | Petrosius V *et al.*, 2023 | 3D | 519<br>(420 / 99) | 1184 | mESC<br>populations | + | 0.828 |
| 2 | (18) | Hu M *et al.*, 2023 | 3D | 135<br>(105 / 30) | 763 | HEK-293T<br>vs. K562 cells | o | 1.0 |
| 3 | (15) | Krull KK *et al.*, 2024 | 4D | 143<br>(115 / 28) | 1477 | U-2 OS cells<br>+/- IFN-γ | ++ | 0.745 |

Note: tSNE overlap annotation: o: almost complete separation, +: overlap but cluster formation, ++: almost complete overlap

**Figure 3:**
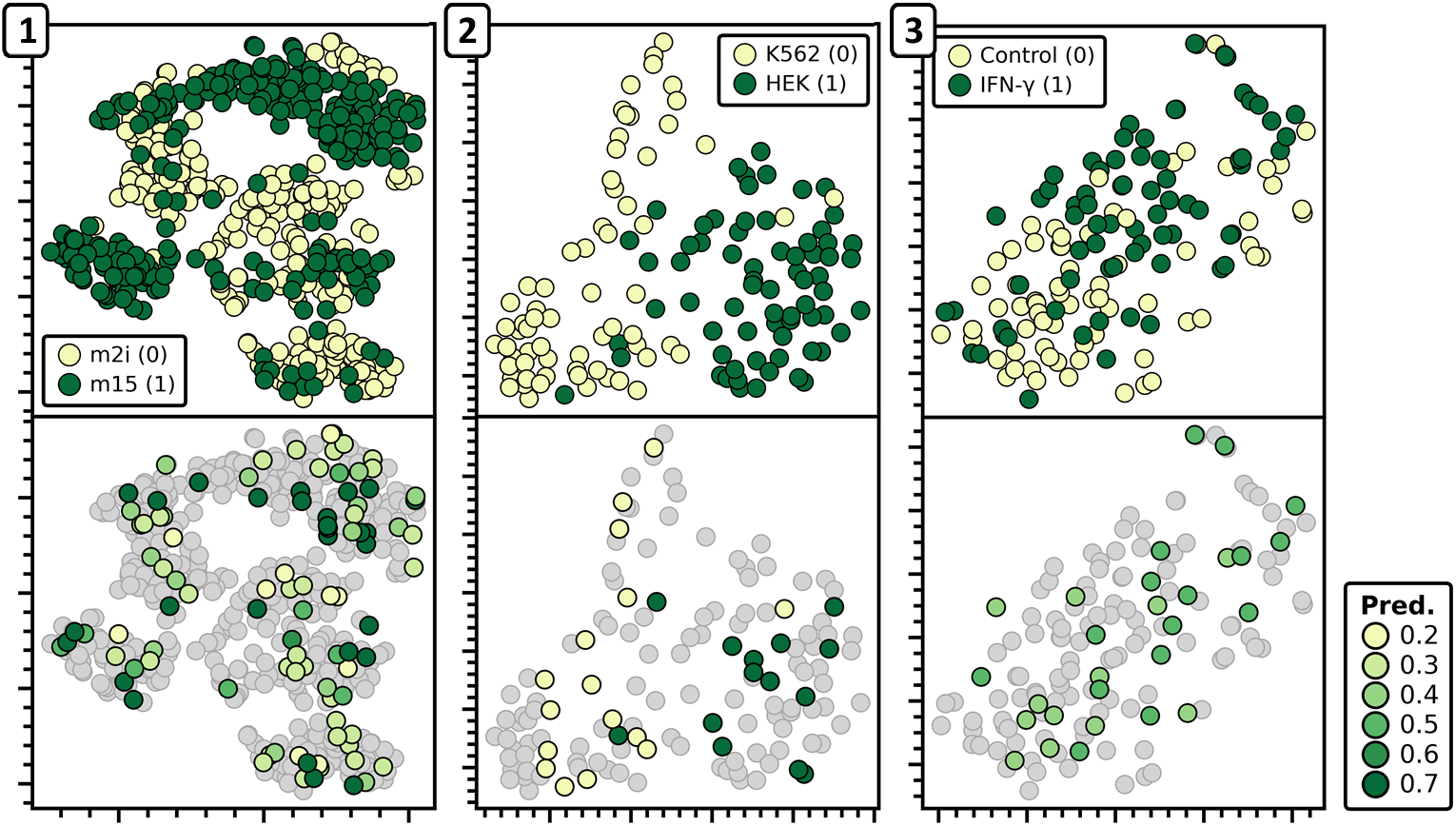
Accuracy of Proformer-based classification of single-cell samples is dependent on quantitative differences. Peptide-level tSNE analysis of four recently published single-cell studies (top panels, numbers refer to Table 1) with varying degrees of quantitative differences between the reported conditions (labels for Proformer training indicated in parentheses). Test set samples with predicated labels after Proformer classification ranging from 0.2 to 0.7 are depicted in the bottom panel (training and validation samples grayed out).

Despite the reduced sensitivity of single-cell measurements that limited the identified peptide number to only around 750 – 1,500 per sample (130,000 – 150,000 in our bulk experiment (**Fig. S1B**)), we achieved particularly high classification accuracies for some of the studies. For instance, the Proformer model distinguished HEK-293T from K562 cells with an AUROC of 100%, indicating its ability to differentiate between highly divergent samples (18), and showed an impressive accuracy of 82.8% for mouse embryonic stem cells (mESCs) (**Table 1**). These cells are biologically much closer than distinct cell lines, however display altered states of pluripotency upon cultivation in different media (17), making them particular challenging to discriminate. For a recently conducted study on U-2 OS cells, the accuracy was 74.5%, likely because of the subtle effect of IFN-γ (15). Here, the limited peptide number (data completeness) in the single-cell data may explain the almost 10%-reduced AUROC value in comparison to our bulk experiment (cf **Fig. 2A**), while the large data set for the mESCs might have benefitted the achieved accuracy. Analyzing all tested single-cell studies by peptide-level tSNE (t-distributed Stochastic Neighbor Embedding) revealed that sample’s predicted labels aligned with their actual label in scenarios where differences in cell populations were more pronounced and displayed values closer to 0.5 especially in the overlapping regions (**Fig. 3**). This finding was in line with our observations made before (cf **Fig. 2C**), suggesting that classification accuracy scales with the quantitative similarity of the tested conditions (indicated by the self-estimated column “tSNE overlap” in **Table 1**), making cases with small differences particularly challenging.

Therefore, additional options need to be explored to improve discrimination even for moderate proteome differences. In this sense, the effect of dimensionality (peptide identifications per samples), data completeness and the number of samples used for training remain to be determined. Here, our work pioneers the use of DL-based classification systems to assign individual cells to unambiguous groups without the need to manually analyze their proteome profiles. We show that the use of our established Proformer pipeline already achieves accuracies up to 100% in cases where cells can be separated based on dimensional reduction approaches. This builds a foundation for follow-up studies to investigate other aspects that help to improve the classification accuracy for samples of various entities and difficulty levels.

## Discussion

We here introduced a transformer-based architecture, called Proformer, for the DL classification of proteomic samples according to their multi-dimensional peptide profile obtained from proteomic data. This model utilizes the values of retention time, mass-to-charge and intensity that are conventionally recorded in LC-MS measurements, and also allows the incorporation of ion mobility as a fourth feature dimension, in a tabular approach that learns characteristics of peptide compositions using a self-attention mechanism. Since the proteome can be considered as the closest reflection of the biological phenotype, classifying proteomic samples by DL approaches has the potential to expand the field from a pure discovery perspective to diagnostic applications, e.g. discriminating healthy and diseased tissues. However, despite advances in high-throughput proteomics (19), the speed at which proteomic samples are analyzed after the measurement might vary from hours to days, limiting the clinical relevance of this technique. In this sense, our Proformer model allows accurate and rapid data interpretation, and could therefore substantially speed up decision-making processes. While others have used deep CNNs to classify proteomic samples (4-8), transformers are based on an attention mechanism, suitable for learning relations inside the input, and enabling the model to capture contextual information effectively without requiring deep architectures. This is particularly helpful for high-dimensional data, as it saves computational costs and makes the model less prone to overfitting. For example, in our bulk data this resulted in almost 50,000-x lower storage allocation for the Proformer compared to the CNNs (data not shown). Moreover, using a tabular transformer approach, we circumvented the binning of data as required in image pipelines (4-8), which, without knowing data relationships, may cause loss of information and protracted data pre-processing. Notably, the Proformer also aggregates the input in its initial step, however, makes use of the learning capabilities of the patch merger to represent the data in an uncorrupted fashion. Given the dimensionality of proteomic data, it is crucial to provide any DL model with sufficient information that reflects the underlying biology in an optimal manner, and we therefore constructed and evaluated our Proformer model using an in-house generated data set that comprised a total of 213 in-depth analyzed samples (**Fig. 1**). Conditions of this experiment came from an IFN-γ treatment of HeLa cells, anticipating that this exposure would only lead to a mild effect in the proteome (14, 15). Indeed, quantitative peptide data of both conditions showed incomplete separation (**Fig. 1E**), therefore allowing optimization of our model, and not reaching perfect classification on the first attempt. As data of the test set were collected long after the training set, we ensured independent evaluation of our models, and found that the Proformer was able to generalize the classification problem, while the CNNs showed a substantial drop in their performance in comparison to the validation set (**Fig. 2A** and **B**). This constitutes an essential finding as it suggests that post hoc measurement of test samples can be classified by the Proformer irrespective of the time and instrument calibration, and potentially even other limiting factors, such as instrument types. As an additional feature dimension for our tabular models, we further proposed the inclusion of ion mobility, allowing the separation of isobaric peptides. We demonstrated its significant impact on the decision process, contributing to a higher accuracy for our 4D-Proformer in comparison to only three features (**Fig. 2A** and **D**). Independent of the model, the data in this work profited from the use of ion mobility as it filters contaminating ions during the measurement, which conveniently leads to cleaner spectra and more peptide identifications (9). Still, among all used dimensions, ion mobility has the lowest resolution (around 50 at typical 100 ms ramp time (20)), offering potential for future optimization, which possibly further refines the classification accuracy. Finally, to explain the decisions of the Proformer, we applied the SHAP value system (16), and identified trends of feature dimensions that contributed differently to the model (**Fig. 2E** and **S2A**), while we proposed a connection of these findings with the quantitative variability of the respective peptides (**Fig. S2B**). Moreover, when we applied the Proformer to a number of publicly available single-cell data sets, we found a relationship between quantitative differences of experimental conditions and the performance of the classification task, which we could also observe in our bulk data (cf **Fig. 2C, Fig. 3** and **Table 1**). While this observation might not be surprising, it testifies that decisions reflect biological differences in the peptide profiles, and encourages optimization to improve accuracy, e.g. by the model design or number of addressed training samples. In this sense, the Proformer should provide an innovative approach for the accurate classification of proteomic samples of different entities that were shown in this work, and might be a pivotal step towards transition protein analysis to clinical applications.

## Material & Methods

### Cultivation of HeLa cells and Interferon-gamma treatment

Human HeLa S3 cells were cultivated in DMEM medium (Gibco), supplemented with 10% (v/v) fetal bovine serum (FBS, Gibco) and 2 mM GlutaMAX (Gibco) using six 10-cm culture dishes. As cells reached 70% confluence, FBS concentration was reduced to 3% (v/v) and cultures were incubated for another 1 hour until three dishes received treatment with 100 ng/mL recombinant IFN-γ (Cell Signaling, diluted in phosphate-buffered saline (PBS, Sigma-Aldrich)) for 24 hours.

After incubation, the culture medium was aspirated, and cells underwent washing by pre-warmed PBS. Adherent cells were detached by gentle scraping in 10 mL PBS, and afterwards sedimented at 1,000 x g for 5 min. The supernatant was discarded and cells were washed once in 5 mL ice-cold PBS followed by centrifugation at 1,000 x g for 5 min and removing the remaining buffer from the pellet. Cells were frozen and stored at −80 °C, preserving their integrity until their following preparation.

### Preparation of proteomic samples

For the preparation of the training and test sample set, obtained pellets of treated and control cells were thawed and lysed in 50 mM TEAB, pH 8.5 (Sigma-Aldrich) buffer + 8 M urea (Bio-Rad) and 1x cOmplete™ protease inhibitor (Sigma-Aldrich). The lysate was sonicated for 15 cycles of 30 sec ON/OFF at 4 °C using a Bioruptor (Diagenode), frozen at −80 °C and afterwards separated from cell debris by centrifugation at 18,000 x g. Samples were transferred to new vials, before protein reduction by adding 10 mM final concentration of dithiothreitol (DTT, Biomol) and incubation at 37 °C and 600 rpm for 45 min. Cysteine residues were alkylated by 55 mM final chloroacetamide (CAA, Sigma-Aldrich) at RT and 600 rpm in the dark for 30 min. Samples were diluted in a 1:6 ratio using digestion buffer (50 mM TEAB + 2 mM CaCl_2_ (Sigma-Aldrich), pH 8.5) to an urea concentration of < 1.6 M and proteins were digested overnight at 37 °C and 800 rpm using sequencing-grade modified trypsin (Promega) at a protein-enzyme-ratio of 25:1. After digestion, peptides were cleaned-up on Sep-Pak cartridges (Waters) containing C_18_ material. The columns were prepared by applying pure ACN (Biosolve Chimie), solvent B (50% (v/v) ACN + 0.1% (v/v) FA (Biosolve Chimie) in ddH_2_O) and solvent A (0.1% (v/v) FA in ddH_2_O), before loading the acidified peptide solution. Samples were washed twice with 1 mL solvent A, and eluted twice in 200 μL solvent B. Purified peptides were frozen at −80 °C, lyophilized in a freeze-dryer and afterwards reconstituted in solvent A. Before samples injection, peptide concentration was determined by BCA assay (Pierce, Thermo Scientific).

### Data acquisition (LC-MS/MS)

In this work, we used a conventional diaPASEF LC-MS methodology that was established in our lab for the measurement of proteomic samples, including fragmentation (sequencing) of peptides to infer their protein identity, however only used peptide information for our subsequent DL models.

For our IFN-γ experiment, we prepared a total of 142 samples for model training and 71 samples for model testing (control and treatment condition; three biological replicates each) that were measured in around six month time difference. To this end, peptide amounts of 100 μg were injected in 2-μL volumes into an EASY-nLC 1200 system (Thermo Scientific) that was connected to a timsTOF Pro mass spectrometer (Bruker Daltonics) via a nano-flow electrospray ionization (nano-ESI) source (Captive Spray, Bruker Daltonics). Peptides were separated on an analytical column (IonOpticks Aurora Series, 25 cm x 75 μm i.d. + CSI, 1.6 μm C18) using an active gradient of 34 min, ranging from 1.6% (v/v) to 28% (v/v) ACN in ddH_2_O (both Biosolve Chimie) at a flow rate of 300 nL/min. During separation, the analytical column was kept at a temperature of 50 °C in a column oven (Sonation). When peptides arrived at the source, they were ionized at a capillary voltage of 1,500 V, a dry gas flow rate of 3.0 L/min and at a temperature of 180 °C. In the MS, ions were accumulated to an IM constant 1/K_0_ of 1.7 V*s/cm^2^ and sequentially ramped from 1.25 to 0.65 V*s/cm^2^ over 100 ms in a locked duty cycle. Subsequent MS1 scans were acquired for peptide signals between 200 and 1,700 m/z, while only precursors with a mass ratio of 475 to 1,000 m/z were considered for DIA window isolation. Eight DIA scans with window widths of 25 m/z and a height of 0.15 V*s/cm^2^ resulted in a cycle time of 0.95 sec. Precursor fragmentation was induced by IM-dependent collisional energies from 45 eV at 1/K_0_ of 1.3 V*s/cm^2^ to 27 eV at 0.75 V*s/cm^2^.

### Raw data processing

Following the measurement, diaPASEF raw data was processed in DIA-NN 1.8.0 (21) using its library-free analysis involving the *in-silico* digestion of a UniProtKB sequence table (FASTA) from *Homo sapiens* (downloaded in September 2021, containing 20,386 reviewed proteins). We allowed a maximum of two missed cleavage sites per peptide and set the peptide length range to 7 – 50 and charge states to 2 – 4. We activated “MBR” to increase data completeness of the respective data set and let protein groups be formed according to protein names from FASTA, while selecting “any LC (high precision)” for precursor quantification. Precursor outputs were filtered at 1% false discovery rate (FDR). Notably, we processed raw data of training and test in two independent processes using equivalent settings.

For the analysis of single-cell data sets that were acquired in DIA mode, we reduced possible missed cleavage sites to a number of one and, in case of the mESC data, further allowed charge states up to five. Analysis of single-cell DDA data (cell line experiment) was performed in MaxQuant (v2.4.2.0) (22) maintaining its default configuration and with “MBR” activated. As protein sequence table, we used the identical FASTA file derived from *Homo sapiens*. Resulting peptide tables were filtered at 1% FDR.

### Preparation of data sets for deep learning pipelines

After raw data processing, we grouped the obtained report tables by individual samples and extracted respective peptide information about sequence, charge state, indexed retention time, indexed ion mobility and MS1 intensity. Here, the use of indexed dimensions fixed them in a uniform range, allowing similar model training when applying different gradient lengths. To calculate monoisotopic peptide masses, we moreover used the “molecular_weight” function in the Bio.SeqUtils.ProtParam (version 1.8.1) (23) Python module and obtained mass-to-charge ratios by the peptide charge states. For the preparation of single-cell studies, we filtered low-identification samples at a threshold of 200 and 800 peptides per cell for the works of Hu M *et al*. (18) and Petrosius V *et al*. (17), respectively, to increase data consistency. Subsequently, tables containing peptide profiles were submitted to the DL pipelines.

### Tabular Proformer pipeline

Before training, extracted peptide data were processed to generate a uniform input: feature dimensions of retention time, mass-to-charge, intensity and ion mobility were z-score normalized, respectively. This steps ensures that all feature have the same scale and increases comparability between samples, allowing an efficient and stable training. Notably, we did not include column embedding, thus fixed the order of feature columns for training and testing. Since individual samples might contain more than hundred-thousand peptides, but show incomplete overlap, we used a patch merger (24) to reduce this dimensionality of the resulting matrix to smaller, equally sized token sets. In this step, sample matrices are divided into non-overlapping patches, each aggregated into one token based on their global context using a self-learning merge mechanism. An additional CLS token is concatenated to the resulting set of tokens, representing the its aggregated information, before passing the matrix to the transformer. The Proformer architecture consists of *N* transformer layers, based on the architecture proposed by the work of Vaswani *et al*. (10), and a classifier head. Each transformer block includes a multi-head attention, followed by a multi-layer perceptron and a normalization layer with residual connections and drop-outs.

The self-attention mechanism of a transformer allows identifying complex structures by an input mapping function. In doing so, each attention head performs three linear learning projections to obtain so-called query, key and value matrices. To determine the output of the self-attention mechanism, a weighted sum of the value matrix are calculated, where the weights are a dot product of the query and key matrices, allowing the model to observe global dependencies. For our desired classification task, we could solely rely on an encoder block as transformer layer. Passing through the encoder blocks, the CLS token is gradually adjusted and used as input for the subsequent classification head to determine the predicted label of the input matrix. During the training process, we used adamax (25) optimization, applied through a cosine warm-up scheduler (26). This pipeline has been implemented in Python code, which can be find in a github repository (link under “code availability”), including all utilized packages.

### Image pipeline

For the CNN model, peptide data was converted to an image, representing retention time by one axis and mass-to-charge by the other axis. To create a uniform data input, this image was divided along a grid and the resulting boxes were binned by their respective summed intensities, followed by normalizing the resulting values per pixel to a range of 0 to 1. Based on previous observations from J. Cadow *et al*. (4), we used ResNet_v2_50 (27), a CNN with a depth of 50 hidden layers that includes batch normalization before each weight layer and Rectified Linear Unit (ReLU) activation functions. ResNet makes use of the residual learning concept, which includes skip connections to jumps over certain layers, allowing the training of very deep neural networks and thus the detection of complex structures. In addition, J. Cadow *et al*. used a pre-trained network based on the LSVRC-2012-CLS data set (28) for image classification. In pre-trained image-based models, early layers have already learned structures, such as edges, textures and basic shapes. Since generated peptide images do not contain such geometric structures, the classification was also performed on a ResNet_v2_50 without pre-trained weights.

### Model tuning

All implemented models were tuned by Optuna (29), based on a tree-structured estimator sampler (30) that fits a Gaussian mixture model for each hyperparameter setting. While training and test data were recorded independently for the bulk IFN-γ experiment, we had to split external data sets into two groups prior to tuning to ensuring unbiased model evaluation. Using a stratified split, training data was divided into five groups, maintaining equal distribution of both labels (experimental conditions). Since proteomic data set sizes are usually limited, we tuned models by aiming for the highest average accuracy (AUROC) among all five validation sets in five-fold cross validation procedure. Subsequently, models were trained with optimal parameter settings using the entire training set, and we assessed their performance by the independent test data.

### SHAP Values

The SHAP (SHapley Additive exPlanations) method (16), established on the Aumann-Shapley game theory (31) provides a framework for interpreting the output of machine learning models. This method assigns a metric to features of individual data points to estimate their contribution to the model’s prediction. To this end, we used the Gradient Explainer method (32, 33) that approximates SHAP values according to the mathematical gradients of the model output, and by considering exemplary input features of reference data points from the training set. As a result, the gradient vector indicates the change rate of the model output for each feature compared to the baseline input. Their sum can be seen as the cumulative impact, while positive value represent a positive contribution and negative values represent a negative contribution, e.g. induced by noisy data.

### Statistical analyses and data illustration

Following raw data analysis of proteomic samples and running DL models, results were evaluated using Python code (version 3.9.12) in the Jupyter Notebook environment (v6.4.8) (34). We used Pandas (v1.4.2) and NumPy (v1.21.6) modules for data handling, and Matplotlib (v3.5.1) and Seaborn (v0.13.1) for data plotting. Dimensional reduction by PCA and tSNE (35) were performed using Scikit-learn (v1.0.2), while illustrative figures of this work were created in BioRender.

## Code availability

An implementation of the described Proformer and CNN image pipeline can be found as Python code at https://github.com/kuehna/proformer.

## Acknowledgement

This work was funded in part by the Federal Ministry of Education and Research (BMBF) and the Ministry of Science Baden-Württemberg within the framework of the Excellence Strategy of the Federal and State Governments of Germany.

