## Supplementary Figures for "Deep learning-based proteomics enables accurate classification of bulk and single-cell samples"

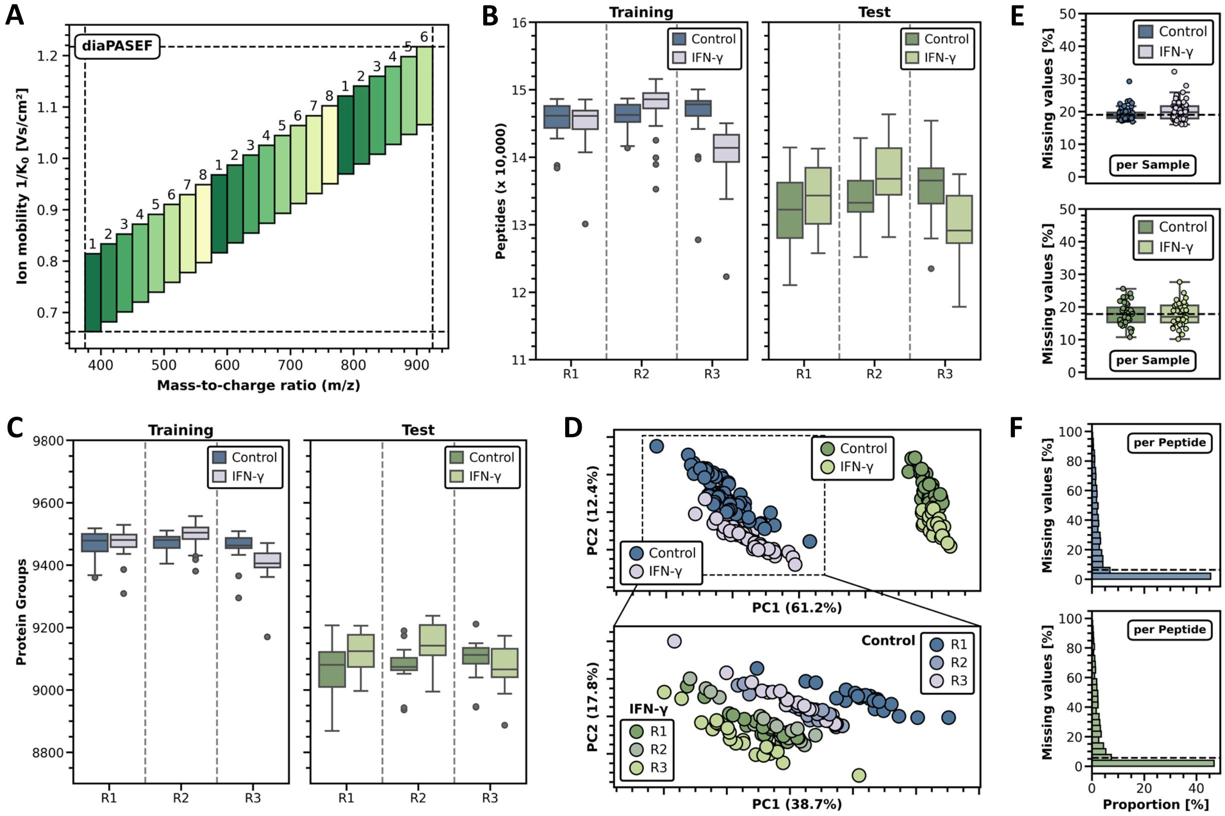


***Supplement Figure S1:*** (A) diaPASEF peptide isolation scheme for subsequent fragmentation. Equivalent window numbers were collectively isolated in one ion mobility ramp. (B) Boxplot distribution of identified peptides per sample and condition in the training (left panel; blue: control, lavender: IFN-γ-treated) and test set (right panel; green: control, light green: IFN-γ-treated). Grey dots indicate outlying samples that exceeded 1.5-x the interquartile range of the distribution represented by the whiskers. (C) Boxplot distribution of Identified protein groups per sample and condition in the training and test set. Same color code and definition as in panel B. (D) Protein-level PCA of training and test set (top panel; same color code than in panel B and C), and enlarged PCA of the training set with indicated replicate numbers (blues: control cultures; greens: IFN-γ-treated cultures). (E) Boxplot distributions of missing value percentages in each of the samples in training (top) and test set (bottom panel). (F) Histogram of missing value percentages per peptide across all samples of training (top) and test set (bottom panel). Bars indicate the share of all peptides.

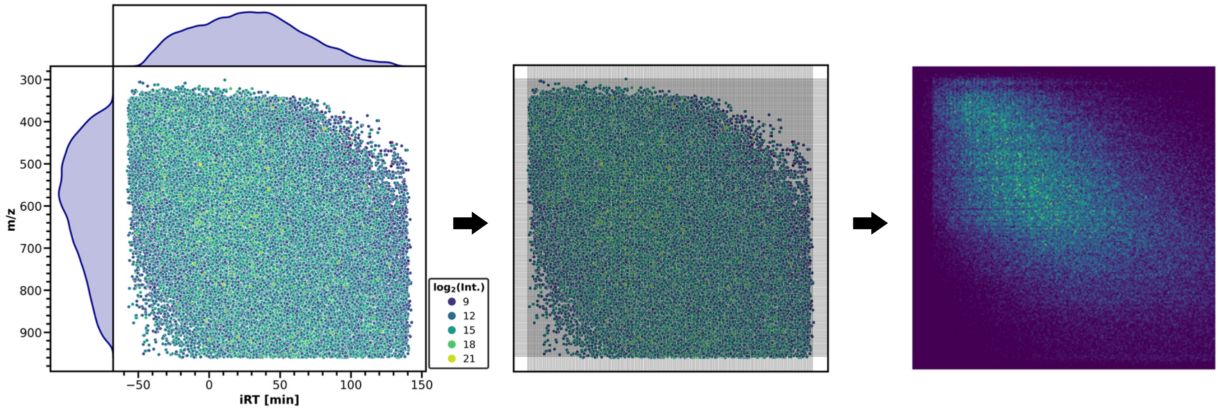


***Supplement Figure S2:*** Image-conversion of LC-MS data using a 224 x 224 grid for CNN pipeline. Left panel: scattered heatmap of peptides in the retention time to m/z space with colors indicating the respective MS1 intensity. Middle: applied grid to bin signals into a 224 x 224 format. Right: resulting heatmap-like image as input for ResNet_v2_50 CNN.

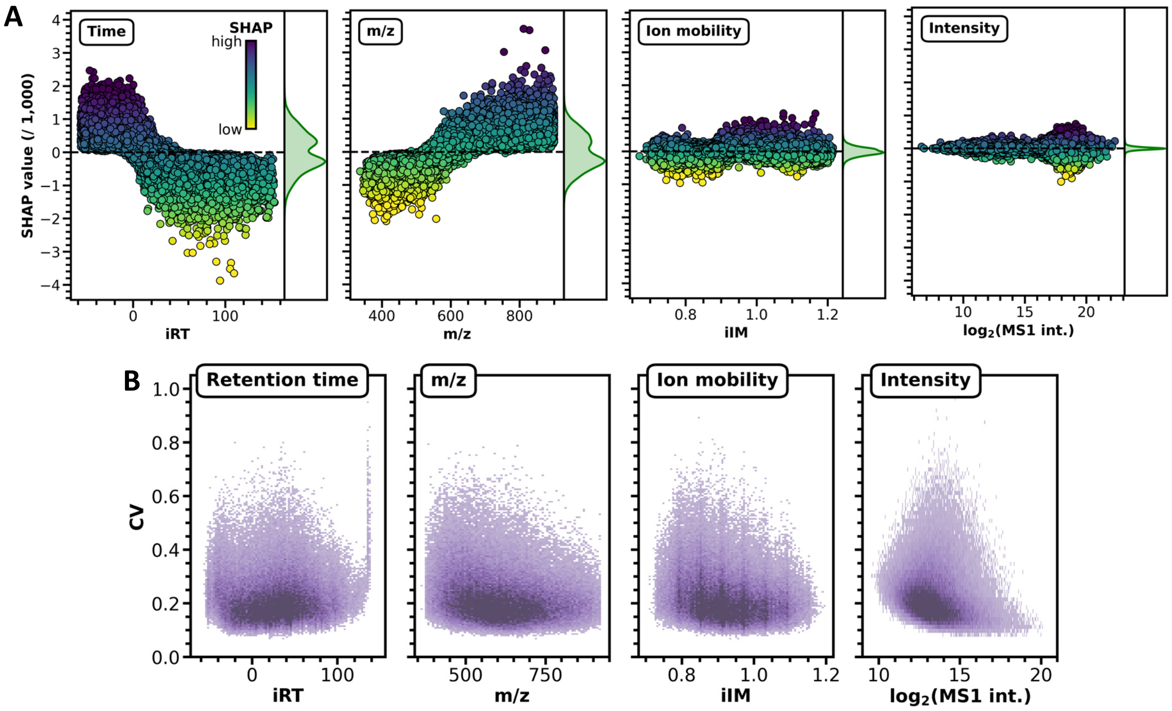


***Supplement Figure S3:*** (A) Distribution of feature-dependent SHAP values in the control condition, showing the contribution of dimension values from individual peptides to the model. High SHAP values (positive contribution) are depicted in blue, while low SHAP values (negative contribution) are shown in yellow. To estimate the density of illustrated data points, SHAP values were additionally shown as kernel distribution. (B) High-resolution 2D-histogram of peptide CV (coefficient of variation) values in the control condition, showing their dependence on the respective feature dimension.
